# Bile Acid Biosynthesis Avoiding Cholesterol

**DOI:** 10.1101/385419

**Authors:** William J. Griffiths, Jonas Abdel-Khalik, Peter J. Crick, Eylan Yutuc, Michael Ogundare, Brian W. Bigger, Andrew A. Morris, Cedric H. Shackleton, Peter T. Clayton, Takashi Iida, Ria Sircar, Hanns-Ulrich Marschall, Jan Sjövall, Ingemar Björkhem, Rajat Rohatgi, Yuqin Wang

**Author notes:** **Corresponding Authors:** Dr William J. Griffiths, Swansea University Medical School, ILS1 Building, Singleton Park, Swansea, SA2 8PP, UK, and Dr Yuqin Wang, Swansea University Medical School, ILS1 Building, Singleton Park, Swansea, SA2 8PP, UK.

## Abstract

Bile acids are the end products of cholesterol metabolism secreted into bile. They are essential for the absorption of lipids and lipid soluble compounds from the intestine. Here we have investigated the bile acid content of plasma and urine from patients with a defect in cholesterol biosynthesis, i.e. Smith-Lemli-Opitz syndrome (SLOS), resulting in elevated levels of 7-dehydrocholesterol (7-DHC), an immediate precursor of cholesterol. Using liquid chromatography – mass spectrometry (LC-MS) we have identified a novel pathway of bile acid biosynthesis in SLOS avoiding cholesterol starting with 7-DHC. This pathway also proceeds to a minor extent in healthy individuals. Monitoring of the pathway products could provide a rapid diagnostic for SLOS while elevated levels of pathway intermediates could be responsible for some of the features of the disease. Importantly, intermediates in the pathway are modulators of the activity of Smoothened, an oncoprotein that mediates Hedgehog signalling during embryogenesis and regeneration of postembryonic tissue.

---

Bile acids are a large family of steroids possessing an acidic group on the side-chain (1). The family can be considered to include both C_24_ acids containing the four-ring steroid skeleton with a 5-carbon side-chain and C_27_ acids with an 8-carbon side-chain attached to the steroid skeleton (2). They are synthesised in the liver (3), but steps in their biosynthesis may also proceed extrahepatically, e.g. in brain (4). Bile acids are synthesised in the liver predominantly via two pathways. The dominating pathway in man is the neutral or normal pathway which starts with 7α-hydroxylation of cholesterol by the hepatic cytochrome P450 (CYP) 7A1 enzyme. The second pathway, known as the acidic pathway, starts with 25R(26)-hydroxylation of cholesterol by CYP27A1 to give (25R)26-hydroxycholesterol ((25R)26-HC) either in the liver or extrahepatically (3,5,6). Note, we use the systematic numbering system to describe (25R)26-hydroxylation of cholesterol according to IUPAC rules, however, much of the literature describes the resulting product as 27-hydroxycholesterol (27-HC) (7). Unless stated otherwise (25R) stereochemistry is generally assumed. Other minor pathways begin with 25-hydroxylation of cholesterol by cholesterol 25-hydroxylase (CH25H) (8) in e.g. activated macrophages (9) or with 24S-hydroxylation of cholesterol by cholesterol 24S-hydroxylase (CYP46A1) in brain (3). Many of the subsequent enzymes converting hydroxycholesterols to bile acids are operative in multiple pathways allowing metabolite crossing between pathways.

The major bile acids in man are cholic, chenodeoxycholic, deoxycholic and lithocholic acids. The latter two are derived from the former two by 7α-dehydroxylation. Ursodeoxycholic is also present in man but rarely as a major bile acid (10). Bile acids are secreted in bile as glycine or taurine conjugates, or in the case of lithocholic acid as a 3-sulfate. Bile acids function in the intestine to aid absorption of lipids, and are recycled to the liver via the enterohepatic system. As well as functioning as detergents in the intestine bile acids are also signalling molecules, regulating their own synthesis via interaction with the farnasoid X receptor (FXR) (11), while intermediates in their biosynthetic pathways from cholesterol are ligands to other nuclear receptors e.g. liver X receptors (LXRs) (12), pregnane X receptor (PXR) (13), vitamin D receptor (VDR) (14), constitutive androstane receptor (CAR) (15) and to G-protein coupled receptors (GPCRs) e.g. EBI2 (16) and TGR5 (17). Interestingly, some steps in bile acid biosynthesis may occur in the nervous system and almost all of the acidic pathway intermediates from cholesterol to bile acids can be found in brain or cerebrospinal fluid (CSF) (4) and many of these intermediates can cross the blood brain barrier providing traffic in and out of the central nervous system (CNS) (18). Cholic acid has been identified in brain (19) and shown to act as a ligand to LXRs regulating the neurogenesis of red nucleus neurons (20), while the C_27_ bile acid 3β,7α-dihydroxycholest-5-en-(25R)26-oic acid (3β,7α-diHCA) has been shown to regulate the survival of motor neurons, again through interaction with LXRs (21).

It is now accepted that bile acid biosynthesis does not only provide detergent molecules essential in the intestine but also numerous signalling molecules important in a diverse array of biological processes. Unsurprisingly, deficiency in enzymes of the bile acid biosynthesis pathways lead to disease (22), however, as a consequence of the redundancy provided by multiple pathways, often not to a total elimination of bile acid formation e.g. in cerebrotendinous xanthomatosis (CTX) where there is a deficiency of CYP27A1 but some cholic acid formation is maintained (23). Likewise defects in cholesterol biosynthesis result in clinical disorders (24), however, it is unknown if there is sufficient metabolic redundancy for cholesterol to be bypassed and bile acid biosynthesis still maintained by yet another metabolic pathway. In the current work we show how bile acids can be biosynthesised from 7-dehydrocholesterol (7-DHC), an immediate precursor of cholesterol, in patients with the disorder Smith-Lemli-Opitz syndrome (SLOS) where the enzyme 7-dehydrocholesterol reductase (DHCR7) is deficient and 7-DHC is abundant in tissues and plasma (Figure 1). The biological significance of this pathway from 7-DHC is discussed in light of the recent observation that pathway intermediates bind to and activate the extracellular cysteine rich domain (CRD) of the oncoprotein Smoothend (Smo), a GPCR which is involved in Hedgehog (Hh) signalling during development (25) and our finding that 3β-hydroxy-7-oxocholest-5-en-(25R)26-oic acid (3βH,7O-CA), another pathway intermediate, binds to Smo and inhibits Hh signalling. Many of the malformations found in SLOS are consistent with impaired Hh signalling (26).

**Figure 1.**
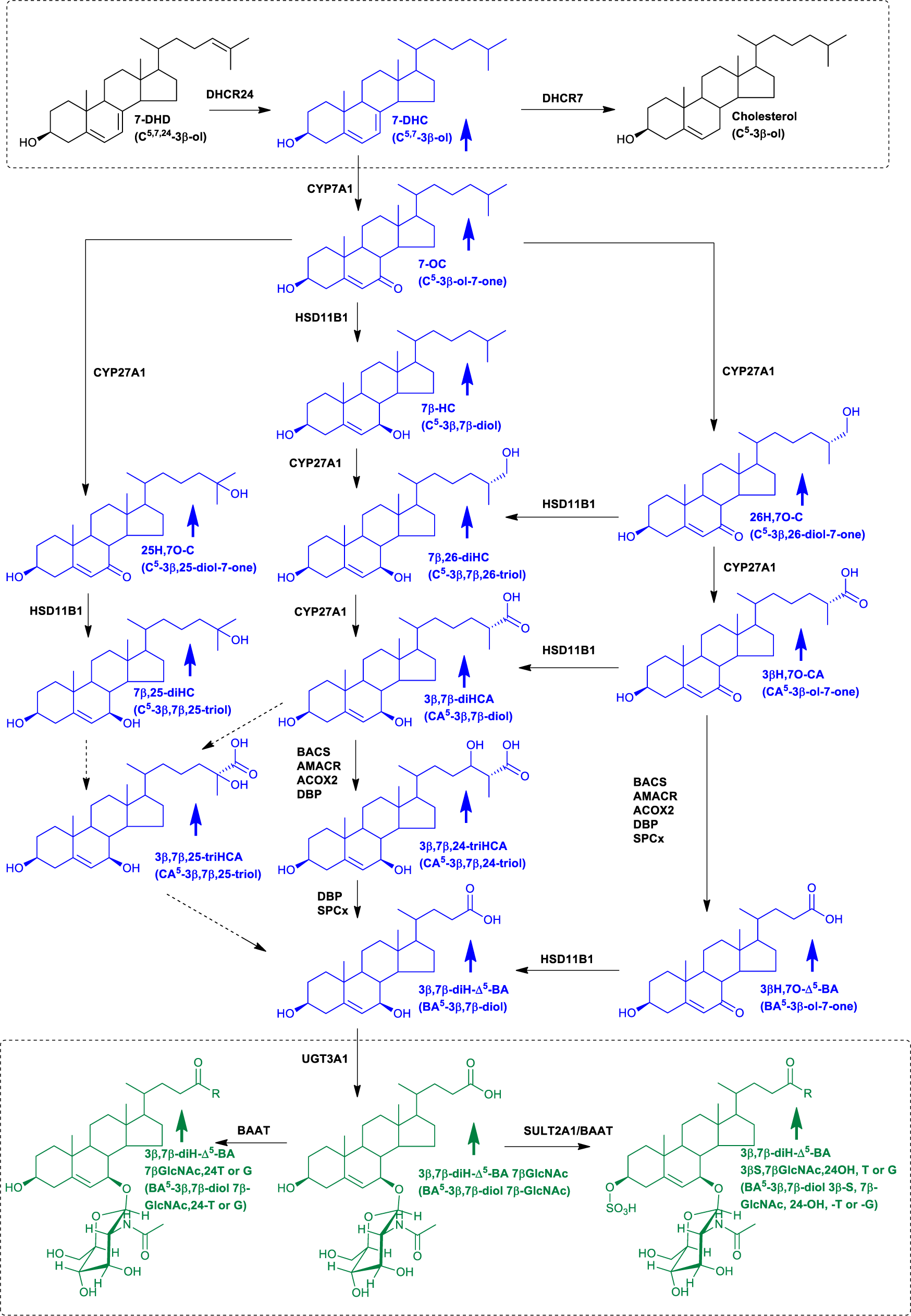
Bile acid biosynthesis starting with 7-DHC and ending with GlcNAc conjugates of 3β,7β-diH-Δ^5^-BA. The metabolites of elevated abundance found in plasma from SLOS patients are indicated by an upward pointing blue arrow. Metabolites of elevated abundance found in SLOS urine are indicated by a green upwards pointing arrow. Oxysterols, including C_24_ and C_27_ acids, found in plasma are shown in blue, bile acid conjugates found in urine are in green. The enzymes for each step are in black. Enzymes, DHCR24, 24-dehydrocholesterol reductase; DHCR7, 7-dehydrocholesterol reductase; CYP7A1, cytochrome P450 family 7 subfamily A member 1; CYP27A1, cytochrome P450 family 27 subfamily A member 1; HSD11B1, hydroxysteroid (11-beta) dehydrogenase 1; BACS, bile acyl CoA-synthetase; AMACR, alpha-methylacyl-CoA-racemase; ACOX2, peroxisomal acyl-coenzyme A oxidase 2; DBP, peroxisomal multifunctional enzyme type 2 or HSD17B4; SPCx, sterol carrier protein x; UGT3A1, 7β-hydroxy bile acid UDP *N*-acetylglucosaminyl transferase; SULT2A1, sulfotransferase family 2A member 1; BAAT, bile acid-CoA:amino acid *N*-acyltransferase.

## Results

### Plasma Analysis

It has recently been shown that CYP7A1 can convert 7-DHC to 7-oxocholesterol (7-OC) (27) and it is known that 7-OC can be converted to 7β-hydroxycholesterol (7β-HC) by hydroxysteroid dehydrogenase (HSD) 11B1 in man and rodents (Figure 1) (28,29). This suggested to us that patients suffering from SLOS may use 7-DHC as a starting point for bile acid biosynthesis rather than cholesterol. We thus investigated using LC-MS exploiting a charge-tagging approach (see Supplemental Figure S1) (30) the nature of bile acid intermediates found in plasma from patients suffering from SLOS. We were able to detect elevated levels of 7-DHC and its metabolites, 7-OC and 7β-HC (reported in (31)), and here we report elevated concentrations of the down-stream metabolites 3β,7β-dihydroxycholest-5-en-26-oic (3β,7β-diHCA) and 3β,7β-dihydroxychol-5-en-24-oic (3β,7β-diH-Δ^5^-BA) acids in SLOS plasma (Figure 2 & 3, Supplemental Table S1, Supplemental Figure S2). Note, we did not include a saponification step in our sample preparation protocol, so concentrations of C_27_ sterols refer to non-esterified molecules. This is justified as it is the non-esterified molecules that have been identified to be biologically active. In some patient samples where the 7-DHC + 8-DHC to cholesterol ratio is high (7-DHC isomerises to 8-DHC (31)), 3β,26-dihydroxycholest-5-en-7-one (26-hydroxy-7-oxocholesterol, 26H,7O-C), 7β,26-dihydroxycholesterol (7β,26-diHC), 3βH,7O-CA and 3β-hydroxy-7-oxochol-5-en-24-oic (3βH,7O-Δ^5^-BA) acids were also observed (Figure 1–3, Supplemental Table S1, Supplemental Figure S2). In the absence of an authentic standard the latter compound was identified based on exact mass, MS^n^ spectra and retention time. Other compounds found to be elevated in some of the SLOS samples where the 7-DHC + 8-DHC to cholesterol ratio is high were 3β,25-dihydroxycholest-5-en-7-one (25-hydroxy-7-oxocholesterol, 25H,7O-C) and 7β,25-dihydroxycholesterol (7β,25-diHC) (Supplemental Table S1). Low levels of metabolites with retention time and fragmentation patterns consistent with 3β,7β,24-trihydroxycholest-5-en-26-oic (3β,7β,24-triHCA) and 3β,7β,25-trihydroxycholest-5-en-26-oic (3β,7β,25-triHCA) structures were also presumptively identified by comparison to the 7α-epimers which were available as authentic standards (Figure 3, Supplemental Table S1, Supplemental Figure S2). These twelve metabolites fall on three branches in pathway from 7-DHC to 3β,7β-Δ^5^-BA (Figure 1).

**Figure 2.**
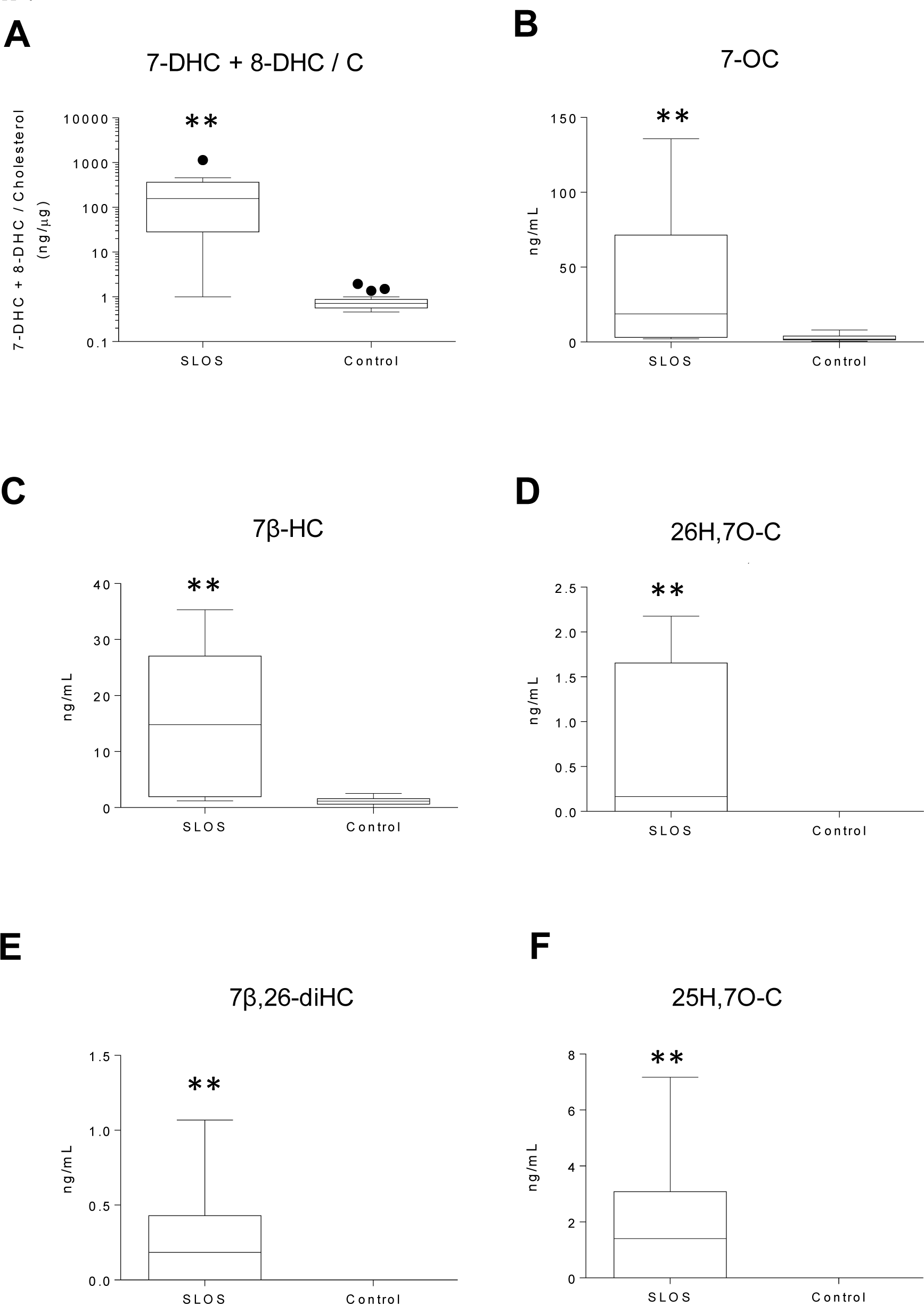
Concentration of 7-DHC and its metabolites in SLOS (n = 10) and control plasma (n = 24). (A) Concentration of 7-DHC plus 8-DHC in ng/µg cholesterol. 8-DHC is an isomerisation product of 7-DHC. Concentrations of all other oxysterols in ng/mL. (B) 7-OC. (C) 7β-HC. (D) 26H,7O-C. (E) 7β,26-diHC. (F) 25H,7O-C. Concentrations determined by LC-MS exploiting charge-tagging utilising the Girard P reagent (30). See also Supplemental Figure S2. The bottom and top of the box are the first and third quartiles, and the band inside the box represents the median. The whiskers extend to the most extreme data points which are no more than 1.5 times the range between first and third quartile distant from the box. Points beyond that are plotted individually. Non-parametric Mann-Whitney test was used for pair-wise comparison for non-normally distributed data. *, P<0.05; **, P<0.01. Data for 7-DHC + 8-DHC, 7-OC and 7β-HC has been reported in (31).

**Figure 3.**
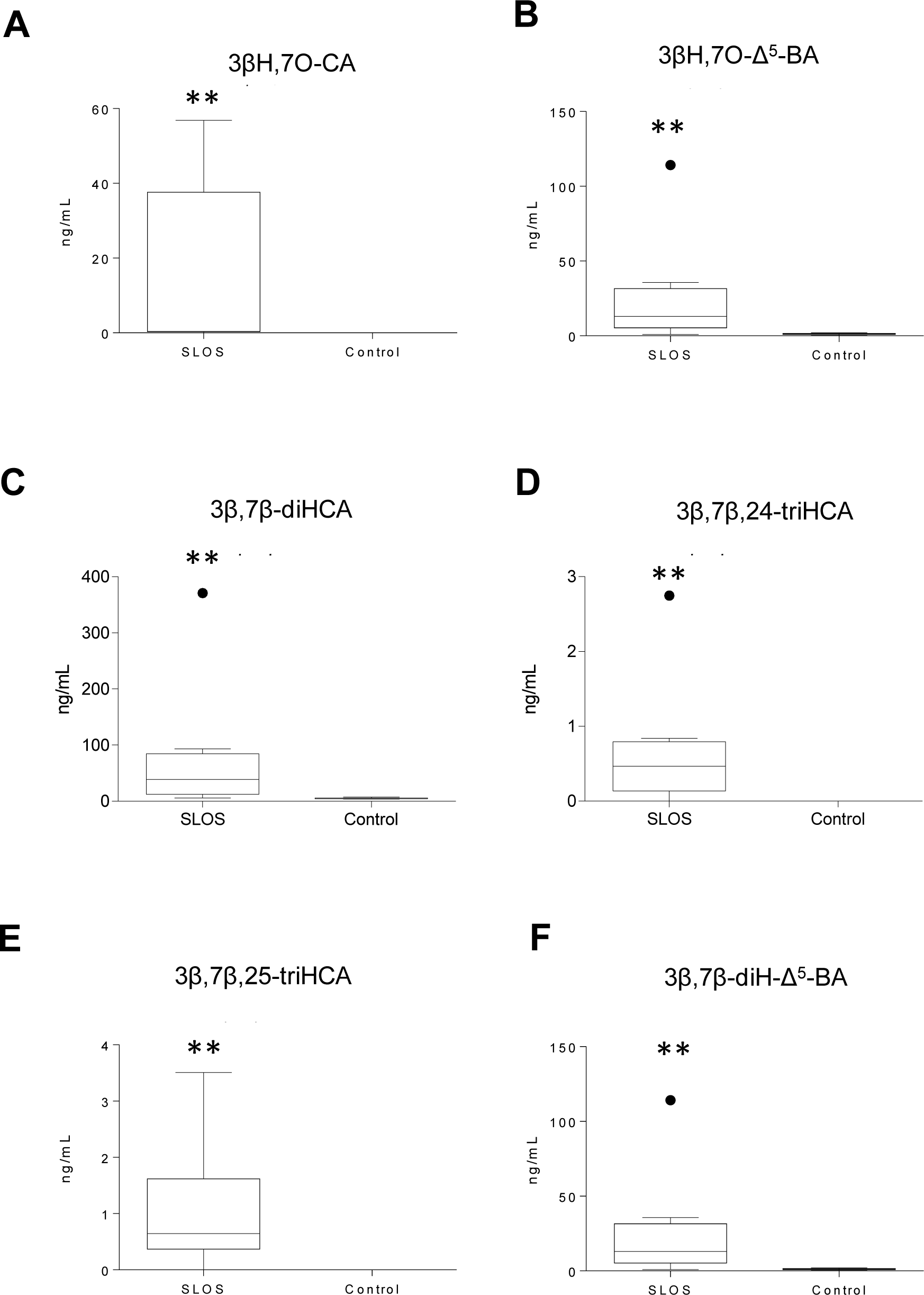
Concentration (ng/mL) of 7β-hydroxy and 7-oxo C_24_ and C_27_ acids in SLOS (n = 10) and control plasma (n = 24). (A) 3βH,7O-CA. (B) 3βH,7O-Δ^5^-BA. (C) 3β,7β-diHCA. (D) 3β,7β,24-triHCA. (E) 3β,7β,25-triHCA. (F) 3β,7β-diH-Δ^5^-BA. See also Supplemental Figure S2. Concentrations determined and statistical comparisons made as described in the caption to Figure 2.

### Urine Analysis

Sterols with a 3β,7β-dihydroxy-5-ene function are not substrates for HSD3B7, the oxidoreductase required to initiate A/B ring transformation ultimately leading to the 3α-hydroxy-5β-hydrogen configuration found in primary bile acids (3,32), so the 3β,7β-dihydroxy-5-ene structure is maintained in the products of the 7-DHC bile acid biosynthesis pathway (Figure 1). Sterols possessing a 7β-hydroxy group are known to be conjugated with *N-*acetylglucosamine (GlcNAc) and excreted in urine (33), hence we next investigated the urine of SLOS patients for GlcNAc conjugated bile acids. We found elevated levels of 3β,7β-diH-Δ^5^-BA conjugated with GlcNAc at position 7β in urine from SLOS patients and also the double conjugate with sulfuric acid (S) at C-3β. Double conjugates with GlcNAc (C-7β) and also glycine or taurine (C-24) and triple conjugates with glycine or taurine and sulfuric acid were also found to be elevated in SLOS urine (Figures 1 & 4, Supplemental Table S2, Supplemental Figure S3). We also found elevated levels of 3βH,7O-Δ^5^-BA conjugated with sulfuric acid in one of the patient samples analysed and increased levels of the double conjugate with sulfuric acid and glycine in two patient samples.

**Figure 4.**
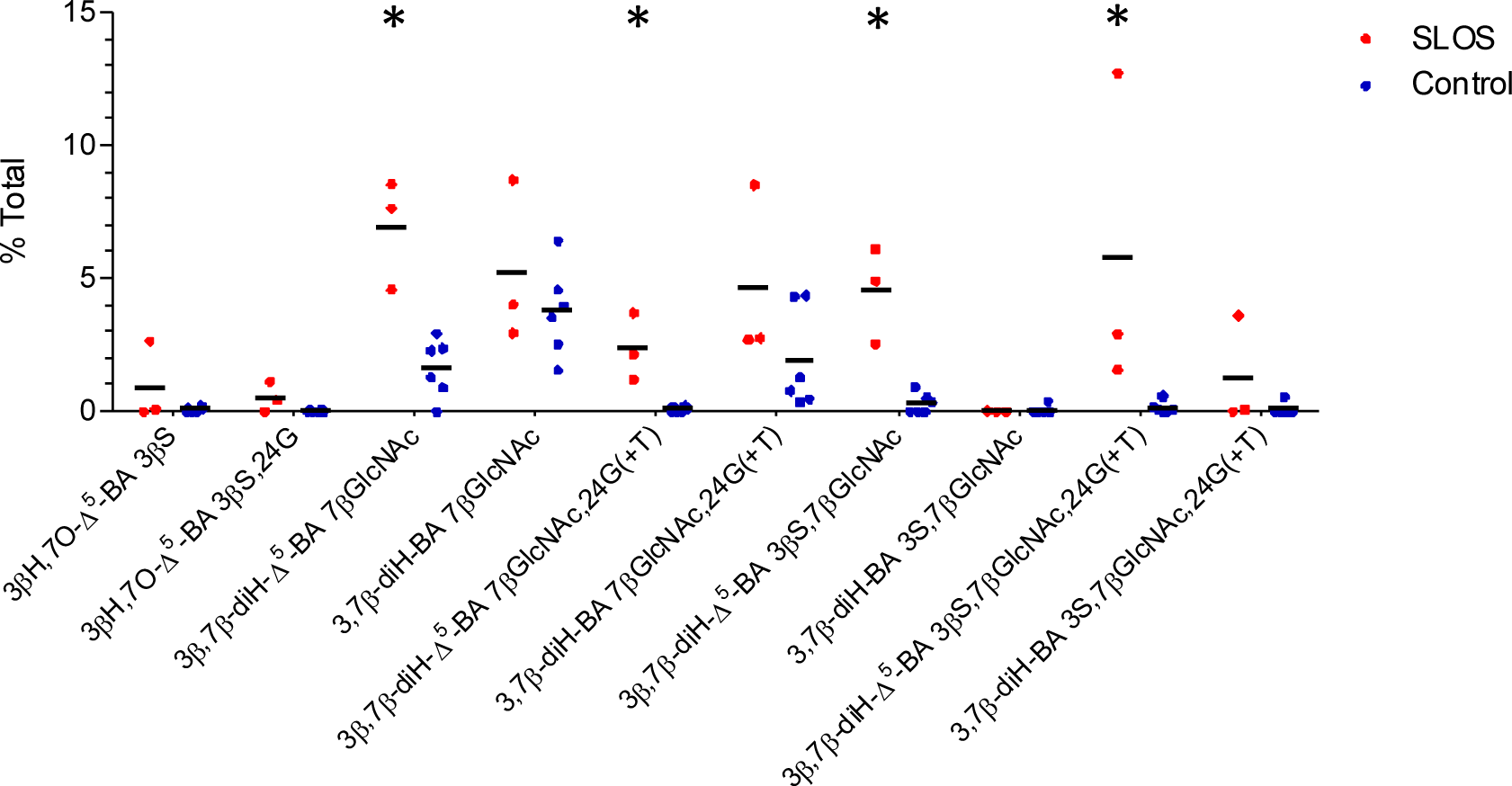
Relative proportions (mole %) of bile acids with 7-oxo or 7β-hydroxy group conjugated with GlcNAc in urine from 3 SLOS patients and 6 controls determined by LC-MS. See also Supplemental Figure S3. Total bile acids include mono-, di-and tri-hydroxylated cholanoic acids and their single and doubly unsaturated equivalents singly, doubly or triply conjugated with sulfuric acid, GlcNAc and glycine or taurine. Non-parametric Mann-Whitney test was used for pair-wise comparison for non-normally distributed data. *, P<0.05; **, P<0.01.

### Hedgehog Signalling

Recent studies have indicated that 25H,7O-C and 26H,7O-C can activate the Hh signalling pathway (25). We speculated that 3βH,7O-CA the downstream metabolite of 26H,7O-C (Figure 1) may also activate the Hh pathway. As reported (25), we also found that both 25H,7O-C and 26H,7O-C induced *Gli1* mRNA, a metric of Hh signalling, in the absence of sonic hedgehog (SHH) protein, but 3βH,7O-CA had the reverse effect and inhibited Hh signalling in the presence of SHH (Figure 5). To investigate if the 7-oxo compounds bind to Smo through its extracellular CRD we used the purified CRD from zebrafish Smo and tested it for binding to 20(S)-HC immobilised on beads in the presence of the various 7-oxo competitors. 20(S)-HC is known to bind to the CRD of vertebrate Smo and to activate Hh signalling (34,35). In the absence of competitor (none) Smo robustly binds to 20S-HC beads (Figure 5) (35). In agreement with an earlier report 26H,7O-C and 25H,7O-C are very good competitors for Smo binding (25), similar 20S-HC. We now find that 3βH,7O-CA also acts as a competitor for the binding of Smo CRD to 20S-beads (Figure 5). These data provide support for the concept that the 7-oxo compounds can bind to the extracellular CRD of Smo and modulate Hh signalling. As SLOS is a disease which phenocopies deficient Hh signalling (26), dysregulation of the balance between the 7-oxo compounds may be an explanation for this deficiency and consequently of malformations seen in SLOS.

**Figure 5.**
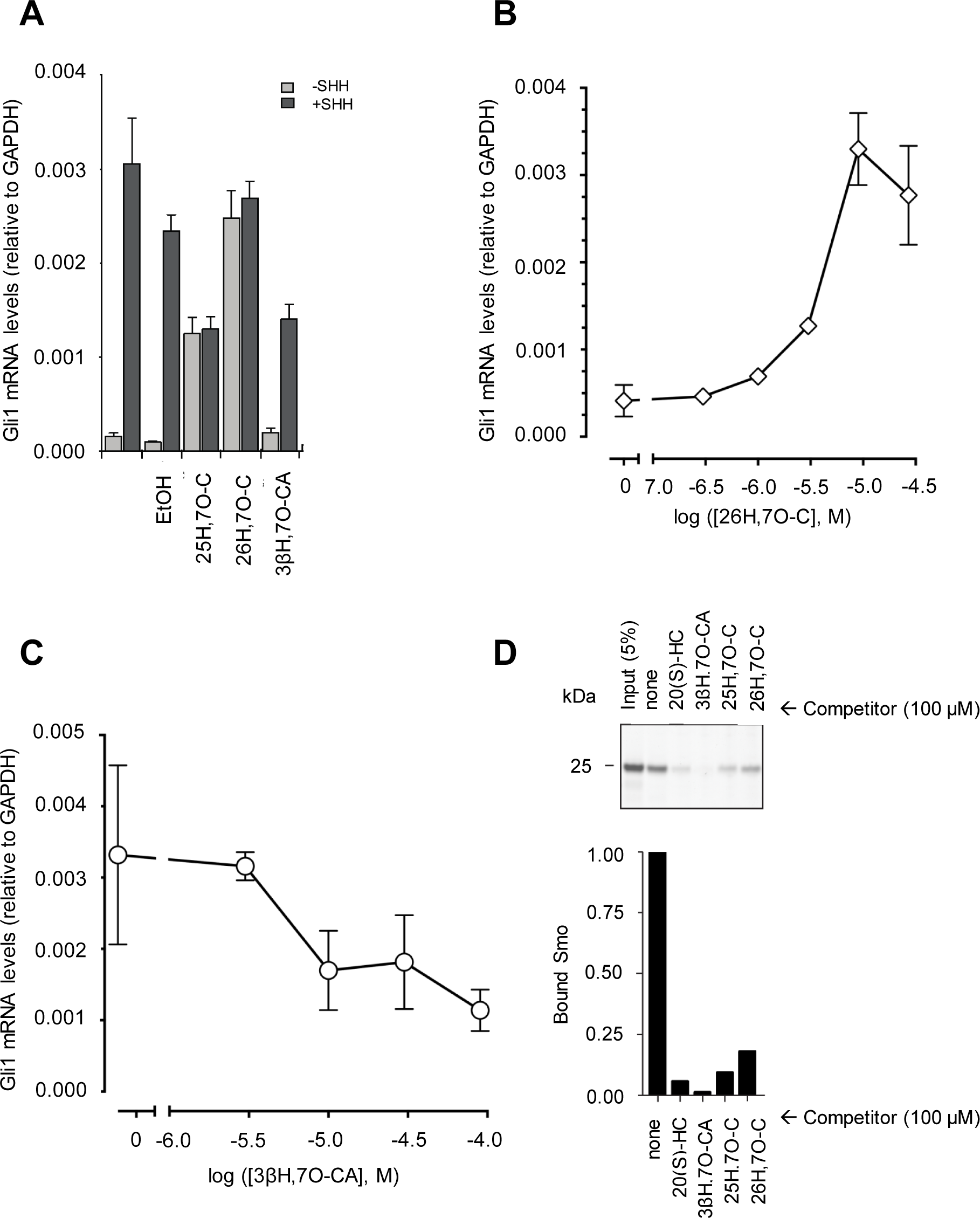
Hh signalling is modulated by 7-oxo metabolites of 7-DHC which bind to the CRD of vertebrate Smo. Levels of *Gli1* mRNA (mean arbitrary units ± SD, n = 4) were used as a metric for Hh signalling activity in NIH/3T3 cells after stimulation (8 hr) with or without 300 nM SHH or 7-oxo compound. (A) Response to 10 µM 7-oxo compound. Dose response curves for (B) 26H,7O-C without SHH and (C) 3βH,7O-CA with SHH. (D) Immunoblots showing the amount of zSmo CRD captured on 20(S)-HC beads in the presence of the indicated competitors.

## Discussion

Bile acid biosynthesis normally starts from cholesterol, however, CYP7A1 can also use 7-DHC as a substrate giving 7-OC as a product (27) which can be reduced by HSD11B1 to 7β-HC (28,29) opening a new route to bile acid biosynthesis (Figure 1). The elevated levels of these sterols in plasma of SLOS patients (31) and also those of 3β,7β-diHCA and 3β,7β-diH-Δ^5^-BA define a new and unexpected pathway for bile acid biosynthesis in SLOS patients (Figures 1 -3). Further evidence for this pathway is provided by the presumptive identification of 3β,7β,24-triHCA, a necessary intermediate as the CoA thioester in peroxisomal side-chain shortening of 3β,7β-diHCA to 3β,7β-diH-Δ^5^-BA (Figure 1). A second branch to the pathway is defined by the identification of 26H,7O-C, 3βH,7O-CA and 3βH,7O-Δ^5^-BA in SLOS plasma from patients with a high 7-DHC + 8-DHC to cholesterol ratio, each of which presumably act as a substrate for HSD11B1. A third branch to the pathway may proceed through 25H,7O-C, 7β,25-diHC and 3β,7β,25-triHCA, although it is not known whether CYP27A1 is responsible for the oxidation of the terminal carbon to the carboxylic acid and how the resulting triol undergoes side-chain shortening (Figure 1). The 7β-hydroxy group in bile acids is known to become conjugated with GlcNAc, leading to the excretion of GlcNAc conjugates in urine (33). Screening for bile acids in urine of SLOS patients revealed elevated levels of 3β,7β-dihydroxychol-5-enoic acids conjugated with GlcNAc at the 7β position and also the double conjugate with glycine or taurine or with sulfuric acid and the triple conjugate with GlcNAc, sulfuric acid and glycine or taurine.

Although in this study we have made no attempt to correlate SLOS severity with 7-oxo, 7β-hydroxy or 7β-GlcNAc metabolite levels the possibility exists that some of these metabolites may be of diagnostic or prognostic value. Furthermore, the identification of GlcNAc or sulfuric acid conjugates in urine could have value in prenatal screening programmes.

The data presented above is supported by a study performed on SLOS urine by Natowicz and Evans in 1994 using fast atom bombardment MS (36). Although in the absence of chromatography or MS^2^ they were unable to fully characterise the metabolites, they were able to identify hydroxyoxocholenoic (H,O-Δ-BA) and dihydroxycholenoic (diH-Δ-BA) acids, their sulfuric acid conjugates and also a dihydroxycholenoic (diH-Δ-BA) acid doubly conjugated with sulfuric acid and an aminohexose sugar (36). Based on our studies these bile acid structures correspond to 3βH,7O-Δ^5^-BA and 3β,7β-diH-Δ^5^-BA and their conjugates.

Shoda et al have suggested that enzymatic mitochondrial oxidoreduction of 7α-hydroxy-5-ene sterols to their 7β-hydroxy epimers can occur in liver (37). In fact they found that human liver mitochondria rapidly convert 7α,26-diHC and 3β,7α-diHCA to their corresponding 7β-hydroxy epimers in an isocitrate dependent manner, perhaps through their 7-oxo intermediates, providing an alternative route to 3β,7β-diH-Δ^5^-BA and ultimately to ursodeoxycholic acid (37). Additionally, 7α-hydroxy epimerisation may be catalysed by the intestinal flora during the enterohepatic circulation to produce ursodeoxycholic and ursocholic acids present in bile and urine of healthy humans (33,38). It is likely, that there are several pathways to 7β-hydroxy bile acids in man of possibly varying importance in health and disease.

It is noteworthy that we have previously identified 3β,7β-diH-Δ^5^-BA singly conjugated with GlcNAc at the C-7β position, double conjugated also at C-3β with sulfuric acid and triply conjugated with glycine or taurine in urine of a patient suffering from Niemann-Pick disease type C (NP-C) (39). NP-C is a rare inherited lipid trafficking disorder which amongst other symptoms presents with learning difficulties and progressive intellectual decline and in this way shows similarity to SLOS. In our earlier study on NP-C we also identified 3βH,7O-Δ^5^-BA as the 3β-sulfate and also conjugated with glycine or taurine and speculated that these bile acid precursors were intermediates in the biosynthesis of 3β,7β-diH-Δ^5^-BA 7β-GlcNAc conjugates (39). More recent studies by Maekawa et al suggested 3β,7β-diH-Δ^5^-BA conjugated with sulfuric acid at C-3β and GlcNAc at C-7β as a new urinary biomarker for NP-C disease (40). However, Clayton and colleagues have found that some NP-C patients carry the cT361G mutation in *UGT3A1* leading to the amino acid substitution p.C121G and an absence of the activity of the encoded 7β-hydroxy bile acid UDP *N*-acetylglucosaminyl transferase (41). 20% of the Asian and Caucasian populations are homozygous for this mutation, so it can be expected that 20% of SLOS patients will not show GlcNAc conjugates in urine. In the absence of functional 7β-hydroxy bile acid UDP *N*-acetylglucosaminyl transferase it can be expected that sulfuric acid conjugates of 7β-hydroxy and 7-oxo bile acids would be further elevated in urine. In our small study of urine samples, all SLOS patients excreted GlcNAc conjugates, so this hypothesis was not tested. In the original publication by Alvelius et al it was suggested that the 7-oxo-and 7β-hydroxy bile acids observed may be a consequence of extensive lipid peroxidation of cholesterol in NP-C disease (39). This seems likely to be correct as elevated levels of 7-OC and also cholestane-3β,5α,6β-triol derived enzymatically from 5,6-epoxycholesterol, like 7-OC a peroxidation products of cholesterol, are found in plasma of NP-C patients (41). Importantly, cholestane-3β,5α,6β-triol is not elevated in plasma of SLOS patients indicating that different oxidative mechanisms are at play in SLOS and NP-C. Additionally, the absence or presence of elevated levels cholestane-3β,5α,6β-triol or its further metabolites in plasma should allow the simple differentiation of SLOS from NP-C. The presence of elevated concentrations of 7-OC in NP-C plasma, in this case generated through peroxidation mechanisms, should also lead to a pattern of bile acid precursors similar to those presented in Figure 1. This is in fact the case (unpublished data) and supports the bile acid biosynthesis pathway presented in Figure 1.

The SLOS phenotype is very broad; severely affected cases often die *in utero* or soon after birth, whereas mild cases show only minor physical abnormalities and learning and behavioural problems (26). Limb abnormalities are common in SLOS and in addition to physical malformations SLOS patients have impaired cognitive function although normal intelligence is also possible (26). Some SLOS patients present with cholestatic liver disease, as do some patients with the bile acid synthetic defect CTX and larger number of patients with NP-C disease, perhaps due to a lack of primary bile acid dependent bile flow, as in the primary defects of bile acid biosynthesis, but this could also be due to potential toxicity of abnormal bile acids. Dietary cholesterol supplementation is common. Although the primary enzymatic defect in SLOS is well defined, its pathophysiology is not, and it is unlikely that only one mechanism explains the myriad of symptoms. It is tempting to speculate that some of the metabolites of the newly defined bile acid biosynthesis pathway (Figure 1) explain some of SLOS phenotypic features. In fact, 3β,7β-diHCA is toxic towards neurons and may be responsible for some of the neurological symptoms of the disease (21).

Hh signalling is required for embryonic patterning and regeneration of postembryonic tissue and aberrant Hh signalling has been linked to SLOS (25,42). In fact, many developmental malformations attributed to SLOS occur in tissues where Hh signalling is required for development (43). DHCR7, the defective enzyme in SLOS, has been implicated to function as a positive regulator of Hh signalling, while the cause of some of the developmental abnormalities seen in SLOS have been attributed to cholesterol deficiency interfering with normal Hh signalling (42,44). Alternatively, Koide et al have suggested that DHCR7 functions as a negative regulator of Hh signalling and its inhibitory affect is at the level, or downstream, of the oncoprotein Smo (43). Both of these proposals can be accommodated by the model suggested by Myers et al where oxysterols and cholesterol can bind to and modulate Smo at different structural regions (25). Smo is a seven-transmembrane protein with extended extracellular and cytoplasmic termini. Hh pathway activation is initiated by binding of cholesterol-modified Hh protein to its receptor Patched (Ptch) an antiporter protein which releases inhibition of Smo and triggers transcription of Hh target genes via Gli transcription factors (45,46). It has been shown that an extracellular CRD is the site for oxysterol binding to Smo and suggested that oxysterols may stabilise an active Smo conformation (25,35,47). Recently, Byrne et al determined the crystal structure of Smo and found a cholesterol molecule bound to the CRD (48). They proposed that cholesterol functions as an endogenous Smo ligand that occupies the CRD groove and there is now data that cholesterol is sufficient to activate Hh signalling through the CRD site (49,50). 20S-HC has been shown to activate Smo in vitro (35) but its presence in vivo is under debate (51). Two other oxysterols which we identify in the current study, 25H,7O-C and 26H,7O-C, are also activators of Smo (25). 26H,7O-C has been previously identified in extracts of retinal pigment epithelial cells, and has been shown to be generated from 7-OC by CYP27A1 (52) (Figure 1). As 7-OC is derived from 7-DHC by CYP7A1 oxidation, the identification of 26H,7O-C and 25H,7O-C in SLOS plasma lends weight to the hypothesis of Koide et al that DHCR7, which reduces the pool of 7-DHC substrate by metabolism to cholesterol (Figure 1), functions as an inhibitor of Hh signalling at the level of Smo (43). Although we did not detect 25H,7O-C or 26H,7O-C in plasma from control plasma samples the presence of down-stream metabolites in plasma from healthy individuals indicates that the pathway involving their formation is active in man. Like 26H,7O-C, 3βH,7O-CA has been identified in retinal pigment epithelial cells, derived by CYP27A1 oxidation of 7-OC (52). This acid which is present in SLOS plasma and to a minor extent in control samples is structurally similar to 26H,7O-C but as shown here inhibits Hh signalling (Figure 5). The involvement of three 7-oxo metabolites of 7-DHC (25H,7O-C, 26H,7O-C and 3βH,7O-CA) in Hh signalling can explain conflicting results suggesting DHCR7 is both a positive (42) and negative (43) regulator of Hh signalling with 3βH,7O-CA inhibiting Hh signalling but both 25H,7O-C and 26H,7O-C activating Hh signalling. The balance between these three metabolites and also a fourth, 3β,5α-dihydroxycholest-7-en-6-one, derived from 7-DHC and recently shown to inhibit Hh signalling (53), may explain the broad spectrum of the SLOS phenotype. Interestingly, Roberts et al have shown that Ptch becomes a more effective inhibitor of Smo in nearby cells when expressed in cells enriched in 7-DHC, consistent with Ptch antiporter activity mediating secretion of a 7-DHC-derived oxysterol inhibiting Smo via its extracellular domain in neighbouring cells (54). Besides SLOS other disorders of cholesterol biosynthesis and metabolism present with developmental malformations in tissues where embryonic patterning depends on Hh signalling (26). Like SLOS these may also result from an imbalance of 7-oxosterols.

In summary we have identified a new pathway of bile acid biosynthesis starting from 7-DHC rather than cholesterol which is evident in SLOS patients. The pathway is also active in healthy controls. Three pathway intermediates were found to modulate Hh signalling two in a positive manner and the third as an inhibitor. Significantly, SLOS phenocopies deficient Hh signalling.

## Experimental Procedures

### Materials

Sources of materials can be found in (30,31,33,40). Use of human material confirmed to the Declaration of Helsinki and was provided with written informed consent and institutional review board approval. Control samples were from previously studies made in Swansea (55) and were used to evaluate the specificity of 7β-hydroxy and 7-oxo metabolites as biomarkers of SLOS.

### Extraction and Analysis of Sterols and Oxysterols from Plasma

Sterols and oxysterols were extracted from plasma as described in (30). We utilised a charge-tagging protocol to maximise sensitivity for LC-MS analysis (Supplemental Figure S1). Full details are provided in Supplemental Experimental Procedures.

### Extraction and Analysis of Bile Acids from Urine

Working solutions of [2,2,4,4-²H_4_]cholic acid (20 ng/µL), [2,2,4,4-²H_4_]glycochenodeoxycholic acid (20 ng/µL) and [2,2,4,4-²H_4_] taurochenodeoxycholic (20 ng/µL) were prepared in absolute ethanol. 2 μL (40 ng) of each working solution was added to 994 μL of water in a 2 mL microcentrifuge tube. Urine (100 μL, pH 6 -7) was added drop-wise to the 1 mL of water containing deuterated standards. After 10 min ultrasonication the solution was centrifuged at 17,000 g, 4 °C for 30 min and the supernatant retained. An Oasis HLB (60 mg, Waters, Elstree, UK) column was washed with absolute ethanol (4 mL), methanol (4 mL) and conditioned with water (4 mL). The supernatant from above was loaded onto the column and allowed to flow at 0.25 mL/min. After a 3 mL wash with water bile acids were eluted in 4 x 1 mL of methanol. The first two 1 mL fractions (containing bile acids) were combined, diluted to 60% methanol and analysed by LC-MS and LC-MS with multistage fragmentation MS^n^ in an identical fashion to derivatised oxysterols (30) with the exception that bile acid urine analysis was performed in the negative ion mode.

### Statistics

A non-parametric Mann-Whitney test was used for pair-wise comparison for non-normally distributed data. A P value of 0.05 or less was considered statistically significant.

### Hedgehog Signaling Assays Using Quantitative RT-PCR

NIH/3T3 cells were grown to confluency in Dulbecco’s Modified Eagle’s Medium (DMEM) containing 10% Fetal Bovine Serum (FBS, Optima Grade, Atlanta Biologicals, Flowery Branch, GA). Confluent cells were exchanged into 0.5% FBS DMEM for 24 hours to allow ciliogenesis prior to treatment with oxysterols in DMEM containing 0.5% FBS for ~16 hours. SHH protein carrying a C-terminal hexa-histidine tag was expressed in bacteria and purified as described previously (56). The mRNA levels of *Gli1*, a direct Hh target gene commonly used as a metric for signalling strength, were measured using the *Power SYBR Green Cells-To-CT* kit (Thermo Fisher Scientific, Waltham, MA). The primers used are *Gli1* (forward primer: 5’-ccaagccaactttatgtcaggg-3’ and reverse primer: 5’-agcccgcttctttgttaatttga-3’), *Gapdh* (forward primer: 5’-agtggcaaagtggagatt-3’ and reverse primer: 5’-gtggagtcatactggaaca-3’). Transcript levels relative to *Gapdh* were calculated using the Δelta-Ct method. Each qRT-PCR experiment, which was repeated twice, included two biological replicates, each with two technical replicates.

### Oxysterol Ligand Affinity Chromatography

Purified zSmo ectodomain protein (zebrafish Smo CRD expressed in HEK-293T cells) was diluted in 20 mM Tris pH 8.5, 150 mM NaCl, 0.3% octyl-glucoside prior to addition of competitors (100 µM) and 20(S)-HC beads (1:200) (34). Binding was allowed to proceed for 12 hr at 4 °C, and then the resin was washed and captured protein eluted with four washes with SDS with 100 µM DTT. The zSmo ectodomain was measured by colloidal Coomassie staining (34).

## Acknowledgments

This work was supported by the UK Biotechnology and Biological Sciences Research Council (BBSRC, grant numbers BB/I001735/1 and BB/N015932/1 to WJG, BB/L001942/1 to YW), the Swedish Science Council (to IB) and in the US by NIH/NIGMS (GM106078 to RR) and NIH (5R01HD053036 to CHS). Control urine samples were kindly provided by Lisa Bastin, Joint Clinical Research Facility, Swansea University / ABMU Local Health Board. Members of the European Network for Oxysterol Research (ENOR, http://oxysterols.com/) are thanked for informative discussions.

## Footnote

The content is solely the responsibility of the authors and does not necessarily represent the official views of the National Institutes of Health.

The abbreviations used are: SLOS, Smith-Lemli-Opitz syndrome; 7-DHC, 7-dehydrocholesterol; LC-MS, liquid chromatography – mass spectrometry; CYP, cytochrome P450; CYP7A1, cytochrome P450 family 7 subfamily A member 1; CYP27A1, cytochrome P450 family 27 subfamily A member 1; ((25R)26-HC), (25R)26-hydroxycholesterol; 27-HC, 27-hydroxycholesterol; CH25H, cholesterol 25-hydroxylase; CYP46A1, cytochrome P450 family 46 subfamily A member 1; FXR, farnasoid X receptor; LXR, liver X receptor; PXR, pregnane X receptor; VDR, vitamin D receptor; CAR, constitutive androstane receptor; GPCR, G-protein coupled receptor; CSF, cerebrospinal fluid; CNS, central nervous system; 3β,7α-diHCA, 3β,7α-dihydroxycholest-5-en-(25R)26-oic acid; CTX; cerebrotendinous xanthomatosis; DHCR7, 7-dehydrocholesterol reductase; CRD, cysteine rich domain; Smo, Smoothend; Hh, Hedgehog; 3βH,7O-CA, 3β-hydroxy-7-oxocholest-5-en-(25R)26-oic acid; MS^n^, multistage fragmentation; DMEM, Dulbecco’s Modified Eagle’s Medium; FBS, Fetal Bovine Serum; SHH, Sonic hedgehog; 7-OC, 7-oxocholesterol; 7β-HC, 7β-hydroxycholesterol; HSD, hydroxysteroid dehydrogenase; HSD11B1, short chain dehydrogenase/reductase family 26C member 1; 3β,7β-diHCA, 3β,7β-dihydroxycholest-5-en-26-oic; 3β,7β-diH-Δ^5^-BA, 3β,7β-dihydroxychol-5-en-24-oic; 26H,7O-C, 3β,26-dihydroxycholest-5-en-7-one; 7β,26-diHC, 7β,26-dihydroxycholesterol; 3βH,7O-Δ^5^-BA, 3β-hydroxy-7-oxochol-5-en-24-oic; 25H,7O-C, 3β,25-dihydroxycholest-5-en-7-one; 7β,25-diHC, 7β,25-dihydroxycholesterol; 3β,7β,24-triHCA, 3β,7β,24-trihydroxycholest-5-en-26-oic; 3β,7β,25-triHCA, 3β,7β,25-trihydroxycholest-5-en-26-oic; HSD3B7, short Chain dehydrogenase/reductase family 11E, member 3; GlcNAc, *N-*acetylglucosamine; NP-C, Niemann-Pick disease type C; UGT3A1, UDP glycosyltransferase family 3 member A1; Ptch, Patched; 20S-HC, 20S-hydroxycholesterol.

## Author contributions

WJG and YW conceived and coordinated the study and wrote the paper. RR and RS designed, performed and interpreted Hh signalling experiments. JA-K, PC, EY and MO performed all mass spectrometry experiments. All authors contributed to writing the paper, reviewed and interpreted the results and approved the final version of the manuscript.

